# Re-engineering of *CUP1* promoter and Cup2/Ace1 transactivator to convert *Saccharomyces cerevisiae* into a whole-cell eukaryotic biosensor capable of detecting 10 nM of bioavailable copper

**DOI:** 10.1101/2022.04.04.486947

**Authors:** Bojan Žunar, Christine Mosrin, Héléne Bénédetti, Béatrice Vallée

## Abstract

While copper is an essential micronutrient and a technologically indispensable heavy metal, it is toxic at high concentrations, harming the environment and human health. Currently, copper is monitored with costly and low-throughput analytical techniques that do not evaluate bioavailability, a crucial parameter which can be measured only with living cells. We overcame these limitations by building upon yeast *S. cerevisiae*’s native copper response and constructed a promising next-generation eukaryotic whole-cell copper biosensor. We combined a dual-reporter fluorescent system with an engineered *CUP1* promoter and overexpressed Cup2 transactivator, constructing through four iterations a total of 16 variants of the biosensor, with the best one exhibiting a linear range of 10^-8^ to 10^-3^ M of bioavailable copper. Moreover, this variant distinguishes itself by superior specificity, detection limit, and linear range, compared to other currently reported eukaryotic and prokaryotic whole-cell copper biosensors. By re-engineering the transactivator, we altered the system’s sensitivity and growth rate, while assessing the performance of Cup2 with heterologous activation domains. Thus, in addition to presenting the next-generation whole-cell copper biosensor, this work urges for an iterative design of eukaryotic biosensors and paves the way toward higher sensitivity through transactivator engineering.

**Graphical abstract:** 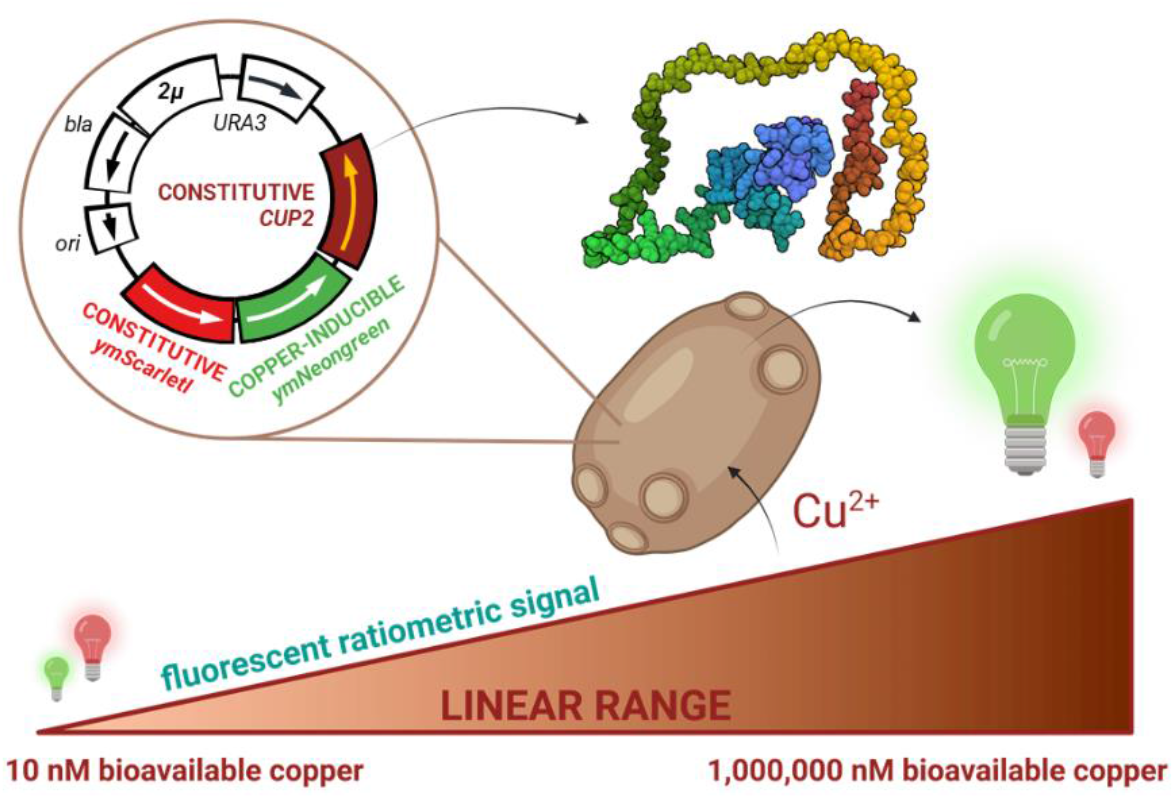

## Introduction

Copper is the 25th most abundant element in Earth’s crust (Emsley, 2001) and, together with iron and aluminium, one of the three most important metals in modern society (Sverdrup et al., 2014). It is indispensable in electronics, transportation, and construction due to its high electrical conductivity, and its popularity keeps rising. For example, each electric car contains over 90 kg of copper, and power plants based on renewable sources need 4-12-fold more copper than those based on fossil fuels. In 2016, 20.2 million tonnes of copper were produced, with a projected annual increase of 6% (Pietrzyk and Tora, 2018). However, copper mining is resource-intensive and polluting, with 70.4 tonnes of water needed to mine one tonne of copper (Northey et al., 2013; Wang et al., 2007).

Copper is also an essential micronutrient but toxic in high concentrations (Valko et al., 2005; Valko et al., 2006). While the lack of copper induces neurological and blood disorders (Madsen and Gitlin, 2007), long-term exposure to high copper concentrations is hepatotoxic (Briffa et al., 2020). On the other hand, copper serves as a fungicide, algaecide, insecticide, and wood preservative (World Health Organization, 2004). Thus, while widely used, copper is closely monitored in drinking water, where it must not rise above 2.0 mg/l (31.5 μM), as water with more than 2.5 mg/l of copper tastes bitter and with more than 4.0 mg/l induces nausea and vomiting (World Health Organization, 2011). Analytical laboratories currently monitor copper using inductively coupled plasma mass spectrometry (ICP-MS), optical emission spectroscopy (ICP-OES), and flame atomic absorption (FAA), thus detecting 0.02 μg/l (0.3 nM) of copper (World Health Organization, 2011). However, these methods are time-consuming, low-throughput, require highly trained staff, and cannot distinguish between total and bioavailable copper.

When assessing pollution, the priority is to quantify bioavailable copper, i.e. copper interacting with living systems (Magrisso et al., 2008), as its amount can differ drastically from that of total copper (Aruoja et al., 2009; Maderova et al., 2011). However, bioavailable copper can be measured only through living organisms, which motivated the development of whole-cell copper biosensors. Such cells are characterised by their specificity (the range of heavy metals they recognise as copper), their detection limit (LOD, the smallest concentration of copper they can detect), operational range (concentrations over which they remain functional), and dynamic range (fold-change in signal strength) (Bhalla et al., 2016).

Most of the current whole-cell copper biosensors were constructed in prokaryotic organisms (Bereza-Malcolm et al., 2015), e.g. *Escherichia coli* (Martinez et al., 2019; Pang et al., 2020), *Pseudomonas* sp. (Li et al., 2014; Maderova et al., 2011), and *Cupriavidus metallidurans* (Chen et al., 2017; Leth et al., 2002), leveraging well-studied bacterial operons induced by heavy metals. However, eukaryotic systems, such as those employing model yeast *Saccharomyces cerevisiae*, might be more relevant to estimate the impact on human health (Walmsley and Keenan, 2000).

*S. cerevisiae* is a non-pathogenic, well-studied, and easy-to-genetically-modify single-cell eukaryote that grows at room temperature. Interestingly, it is highly resistant to copper, a trait probably selected for in vineyards in which copper sulphate was used as a fungicide (Marsit and Dequin, 2015; Steenwyk and Rokas, 2017). This yeast survives high copper concentrations by inducing several genes, including *CUP1* (Shi et al., 2021), which is often present in dozens of tandem repeats (Crosato et al., 2020) and encoding Cu-metallothionein, a well-studied 61-amino-acid protein that binds a surplus of toxic copper and cadmium ions (Ecker et al., 1986).

As it is activated by only a few transcription factors, the promoter of the *CUP1* gene is considered as a model promoter (Ball et al., 2016; Shen et al., 2001) and is often used in chromatin studies (Badi and Barberis, 2002; Mehta et al., 2018; Wimalarathna et al., 2012). Moreover, it is bound by only two transactivators, Cup2 and Hsf1, thus responding only to toxic copper concentrations, heat shock, and oxidative stress (Liu and Thiele, 1996; Tamai et al., 1994). In an environment with too much copper, activation of the *CUP1* gene is mediated by Cup2/Ace1 transactivator. This 225-amino-acid protein detects toxic copper (Singh et al., 2021), activating the transcription of the *CUP1* gene through the C-terminal activation domain after binding copper ions in its N-terminal DNA-binding domain (Li and Fay, 2019; Turner et al., 1998). As such, this transactivator could be employed in biosensor construction, although it remains underinvestigated, with conflicting reports on the feasibility of its overexpression (Shively et al., 2013; Sopko et al., 2006).

In this work, we developed a next-generation eukaryotic whole-cell copper biosensor based on fluorescence monitoring. For this purpose, we designed 16 *S. cerevisiae* plasmids encoding combinations of (i) a dual-reporter fluorescent system, carrying one copper-inducible reporter and another standardising, constitutively-expressed reporter, (ii) four different copper-inducible promoters, and (iii) nine rationally engineered variants of Cup2 transactivator. By this approach, we obtained a strain that detected copper with a linear range from 10^-8^ to 10^-3^ M. By comparing strain performances, we also show that straightforward engineering of transactivators allows construction of highly sensitive eukaryotic whole-cell biosensors.

## Materials and Methods

### Media and growth conditions

*E. coli* was grown overnight at 37°C at either 200 rpm in liquid 2xYT media (16.0 g/l tryptone, 10.0 g/l yeast extract, 5.0 g/l NaCl) or on LB plates (10.0 g/l tryptone, 5.0 g/l yeast extract, 5.0 g/l NaCl, 15.0 g/l agar), supplemented with 100 μg/ml of ampicillin. *S. cerevisiae* was grown at 30°C/180 rpm in chemically defined media without uracil (6.70 g/l Difco Yeast nitrogen base without amino acids, 20.0 g/l glucose, 0.77 g/l MP Bio complete supplement mixture without uracil, and 20.0 g/l agar for solid media). When measuring copper response, Difco’s YNB was exchanged with Formedium’s YNB without amino acids and copper (-ura-Cu medium), to which copper was added in required concentration as CuSO_4_ (Merck, Germany). Biosensor specificity was tested with CoSO_4_, NiCl_2_·6H_2_O, ZnCl_2_, PdCl_2_(NCCH_3_)_2_, Cd(ClO_4_), AgNO_3_, and AuCl (Merck, Germany).

### Plasmid construction

Plasmids were constructed in NEB Stable *E. coli* using Q5 polymerase, PCR Cloning Kit, Q5 Mutagenesis Kit, and HiFi Assembly (New England Biolabs, Evry, France), with details of the constructions given in Supplementary materials. Plasmid DNA was isolated with NucleoSpin Plasmid Mini Kit and extracted from an agarose gel with NucleoSpin Gel and PCR Clean-up Kit (Macherey-Nagel, Duren, Germany). All constructs were verified by restriction digest and Sanger sequencing. Primer synthesis and Sanger sequencing were outsourced to Eurofins Genomics (Konstanz, Germany).

### Yeast strain construction

In this work, all yeast strains were constructed by transforming BY 4742 (Baker Brachmann et al., 1998) as in Gietz and Schiestl (2007). For the flow cytometry analysis, we used strains with overexpression cassette integrated into the chromosome XI at EasyClone locus XI-1 (Jessop-Fabre et al., 2016), at which the neighbouring regions do not interfere with gene expression. This chromosomal insertion ensured that each cell carried only one copy of the expression cassette, thus producing a uniform cell population that could not be obtained with 2μ plasmid due to its varying copy number. Strains with chromosomal insertions were constructed by amplifying overexpression cassettes from previously constructed plasmids with Phusion polymerase (New England Biolabs, Evry, France) and oligonucleotides eo-chrXI-f and eo-chrXI-r (Supplementary materials). BY 4742 was transformed with PCR products and the correct integrations of the transforming DNA fragments, as well as the lack of 2μ plasmids, were confirmed via colony PCR with OneTaq polymerase (New England Biolabs, Evry, France).

### Fluorescence microscopy

Cells were grown overnight in a -ura medium without or with 100 μM CuSO_4_ until they reached a cell density of 10^7^ cells/ml. Next, cells were centrifuged and imaged on a ZEISS LSM 980 confocal microscope with Airyscan 2 (Carl Zeiss, Germany) under a 40x oil objective, using laser wavelengths of 488 and 561 nm. Images were processed using Zeiss Zen Blue software.

### Flow cytometry

Cells were grown overnight in -ura-Cu medium until the cultures reached 5·10^6^ cells/ml and were then supplemented with 100 μM CuSO_4_. Every 30 min, 100 μl of the culture was mixed with 200 μl PBS buffer (pH 7.4), and 25,000 events were recorded on BD LSRFortessa X-20, using 488 nm laser, 530/30 nm emission filter, and 505 nm long-pass dichroic mirror. Each experiment was performed in triplicate. Data were analysed in the R computing environment (R Core Team, 2021) using openCyto (Finak et al., 2014) and ggcyto packages (Van et al., 2018). The gate was computed automatically to enclose 95% of total events on the logicle-transformed forward and side scatter areas. The ymNeongreen was assessed on the flowJo biexponential scale.

### Spectrophotometry

Cells were grown in standard -ura medium to stationary phase, incubated at 4°C for 24 h, washed in sterile deionised H_2_O, diluted tenfold in -ura-Cu medium supplemented with CuSO_4_, and transferred into black sterile Greiner 96-well microplates with a flat bottom. Every well contained 110 μl of mixture and was prepared in triplicate. Microplates were parafilmed to prevent evaporation and shaken at 30°C/180 rpm/16 h after which the parafilm and microplate lid were removed, and the microplate was incubated for 20 min at room temperature.

Next, the microplate was imaged in Clariostar Plus (BMG Labtech, Ortenberg, Germany) preheated to 30°C. The plate was first shaken at 700 rpm/6 min and then imaged for eight cycles with five flashes per well and cycle, with shaking at 700 rpm/30 s between two cycles, after which the average fluorescence value was calculated for each well. The ymNeongreen was excited at 470/15 nm and measured at 517/20 nm, using a 492.2 nm dichroic filter. The ymScarletI was excited at 548/15 nm and measured at 591/20 nm, using a 568.2 nm dichroic filter. Data were exported to .xlsx format and processed in the R programming environment.

### Growth curves

Cells were grown in standard -ura medium to stationary phase, incubated at 4°C for 24 h, washed in sterile deionised H_2_O, diluted tenfold in standard -ura medium and transferred into black sterile Greiner 96-well microplates with a flat transparent bottom so that every well contained 200 μl of the mixture. The microplate, covered with a lid, was put into Clariostar Plus preheated at 30°C and OD_600_ measured in each well every 5 min, for 16 h. Before each measurement cycle, the microplate was shaken at 600 rpm/30 s. Every condition was prepared and measured in triplicate. Data were exported to .xlsx format and processed in the R programming environment.

### Statistical analysis

Statistical analysis was performed in the R programming environment. The coefficient of determination (r^2^) was calculated by fitting a linear model to points within the linear range of the biosensor. Detection limit (LOD) was defined as the minimal Cu^2+^ concentration at which ymNeongreen/ymScarletI ratio was higher than that of cells grown in -ura-Cu medium + its 3 standard deviations.

## Results

We developed a copper biosensor that leverages the native ability of yeast *S. cerevisiae* to detect and detoxify copper ions. The engineered biosensor combines (i) a dual-reporter fluorescence-based system, with one reporter being copper-inducible and another being constitutively-expressed to accurately normalise the signal, (ii) modified *CUP1* promoter and (iii) overexpressed Cup2 transactivator. The constructed biosensor is highly sensitive, detecting as little as 10^-8^ M of bioavailable copper. While constructing it, we also re-engineered Cup2, truncating it and exchanging its native activation domain with heterologous ones, thus highlighting the importance of balanced gene activation.

### The construction strategy

To design an inducible reporter cassette, we first looked at the *CUP1* promoter, which is well-characterised and induced above 10 μM Cu^2+^ (Ecker et al., 1986). Previous studies examined both the original *CUP1* promoter and a hybrid *CUP1-CYC1* promoter (Thiele and Hamer, 1986), mainly focusing on the 150 bp regulatory region upstream of the minimal *CUP1* promoter (Huibregtse et al., 1989) and disregarding an additional 170 bp of regulatory sequence extending to the neighbouring ORF (Figure 1A). While the intensely studied region of 150 bp contains all five Cup2 recognition elements, grouped into distal and proximal upstream activating sequences (UASd and UASp, respectively), the overlooked region also holds putative recognition elements for transcription factors Yox1, Rap1, Eds1, Rlm1, and Sum1 (Castro-Mondragon et al., 2022), which might affect induction. Moreover, the UASd also encodes a heat shock element (HSE) responsive to heat and oxidative stress (Silar et al., 1991), thus promoting non-copper mediated induction. We proceeded by inactivating the unwanted HSE as described previously (Tamai et al., 1994) and combined both *CUP1* and *CYC1* minimal promoters *(minCUP1* and *minCYC1*, respectively) with *CUP1* regulatory regions spanning either 150 bp or 320 bp, thus obtaining four versions of an inducible promoter (Cu1 to Cu4).

**Figure 1:**
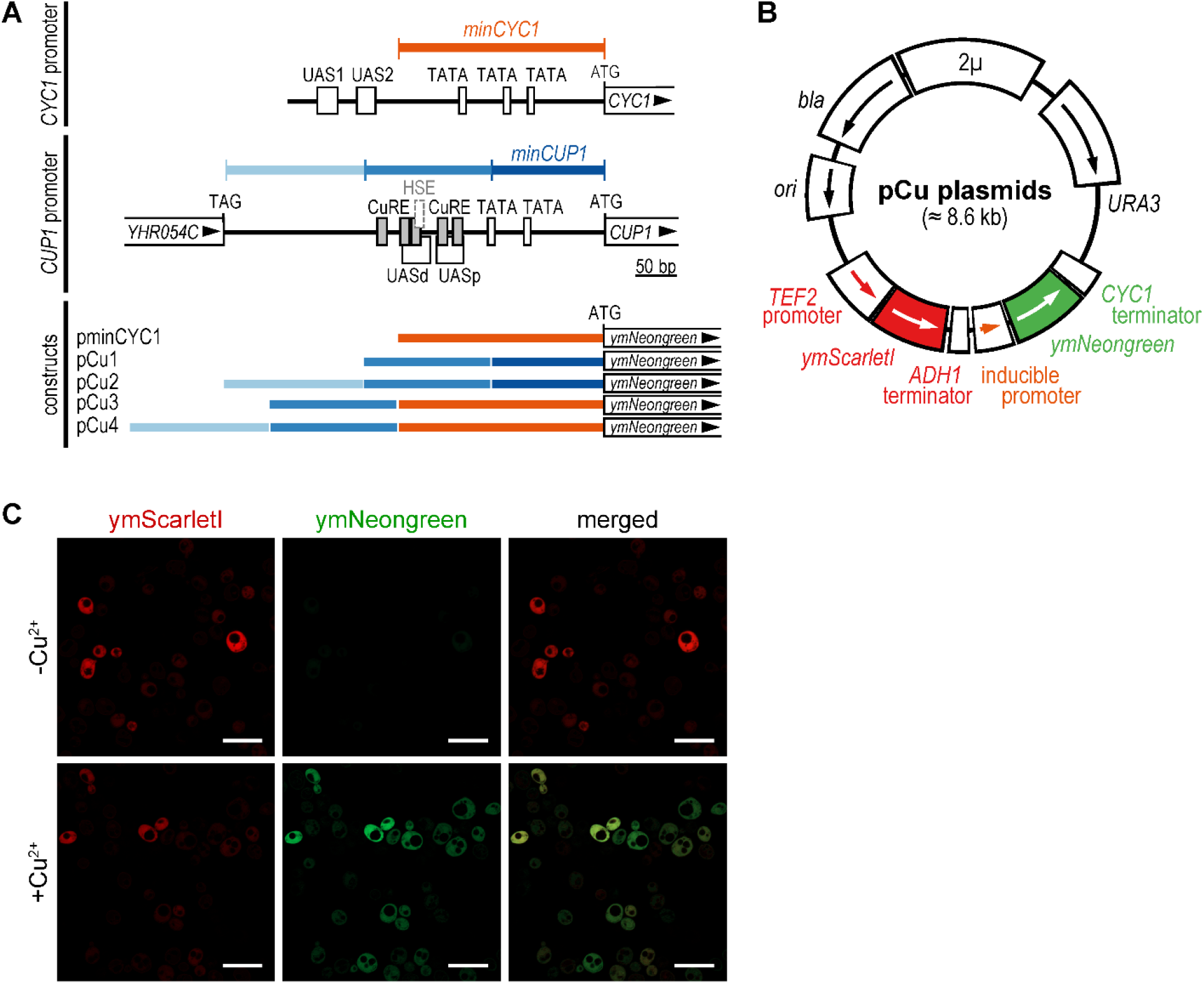
Design of the four copper-inducible promoters. A) Structures of *CYC1* and *CUP1* promoters and four copper-inducible promoters constructed in this work. The *CYC1* promoter has three TATA-boxes within the sequence known as minimal *CYC1* promoter (*minCYC1*) and two upstream activating sequences (*UAS1* and *UAS2*) outside it (Song et al., 2016). The *CUP1* promoter has two TATA-boxes and two upstream activating sequences (UASd and UASp) that encode four Cup2 recognition elements (CuRE) and one heat shock element (HSE) (Thiele and Hamer, 1986). Constructed plasmids carry minimal *CYC1* promoter (pminCYC1), truncated and full *CUP1* promoter (pCu1 and pCu2, respectively) and two hybrid *CUP1-CYC1* promoters (pCu3 and pCu4). B) Plasmids carrying copper biosensors encode β-lactamase *(bla),* pUC19 replication origin (*ori*), constitutive *TEF2* promoter and *ADH1* terminator driving expression of yeast-optimised red fluorescent protein ymScarletI, copper-inducible promoter and *CYC1* terminator driving the expression of yeast-optimised green fluorescent protein ymNeongreen, yeast genetic marker (*URA3*), and replication origin of yeast plasmid 2μ (2μ). C) Fluorescence micrographs of cells carrying pCu2 plasmid, grown in medium without copper (-Cu^2+^) or supplemented with 100 μM copper (+Cu^2+^), expressing ymScarletI and ymNeongreen. Scale bar denotes 10.0 μm.

After constructing inducible promoters, we placed them onto a 2μ plasmid, where they drove the expression of ymNeongreen (Figure 1B). The four constructed plasmids also encoded cassette for constitutive expression of ymScarletI, driven by the *TEF2* promoter. Thus, while ymNeongreen measured Cu^2+^ concentration, ymScarletI reported on cell density, with their ratio indicating Cu^2+^ concentration corrected by the number of cells. Both fluorescent proteins were codon-usage optimised for yeast, displaying high *in vivo* brightness and photostability (Botman et al., 2019) and fast maturation time (Bindels et al., 2017; Shaner et al., 2013). Finally, to ensure a robust signal even from non-induced promoters, we selected *URA3* as a genetic marker, thus maintaining plasmid at over 20 copies per haploid cell (Karim et al., 2013). Fluorescence micrographs showed that ymScarletI was indeed expressed in all conditions, whereas ymNeongreen was expressed only in the presence of Cu^2+^ (Figure 1C), thus demonstrating that the system is fully functional.

### Modifications of the *CUP1* promoter

We then compared our four constructs carrying inducible promoters (pCu plasmids) with the construct carrying only *minCYC1* promoter (pminCYC1). Cells carrying only pminCYC1 exhibited no change in their ymNeongreen/ymScarletI ratio, whatever the Cu^2+^ concentration (Figure 2A). In contrast, cells carrying pCu plasmids increased their fluorescence ratio as Cu^2+^ concentration rose, with their sensitivity and dynamic ratio chiefly determined by their minimal promoter. The constructs built upon *minCUP1* promoter (pCu1 and pCu2) outperformed those built upon *minCYC1* promoter (pCu3 and pCu4), having a higher dynamic range and a detection limit reaching 10^-7^ M Cu^2+^.

**Figure 2:**
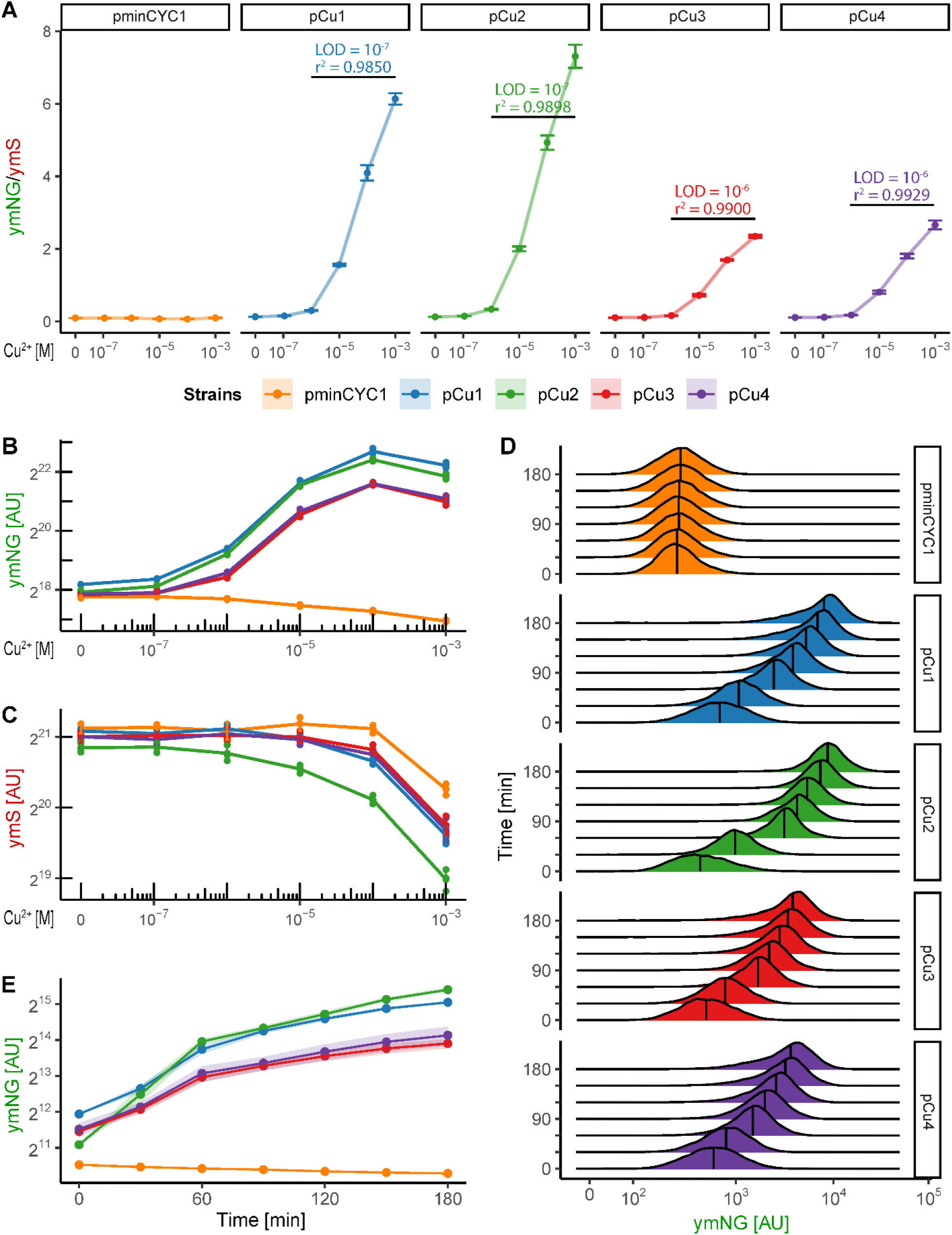
Copper biosensors with different promoters. A) The ymNeongreen/ymScarletI ratio of control construct (pminCYC1) and engineered copper biosensors pCu1-pCu4. LOD = limit of detection, r^2^ = coefficient of determination. Black horizontal lines indicate a linear range. Error bars denote standard deviations (SD). B) ymNeongreen signal. Error bars denote SD, while each point shows one measured sample. C) ymScarletI signal. D) The distribution of ymNeongreen signal in cells induced with 100 μM Cu^2+^, measured by flow cytometry. The ymNeongreen cassette is integrated into the chromosome. Vertical lines within distributions denote median fluorescence. E) Kinetics of promoter induction with 100 μM Cu^2+^ in cells with ymNeongreen cassette integrated into the chromosome, measured by flow cytometry.

A detailed study of the ymNeongreen and ymScarletI signals (Figures 2B and 2C) offered additional insights. The changes in measured ratios originated only from ymNeongreen, except at the Cu^2+^ concentrations higher than 10^-4^ M, at which ymNeongreen plateaued and ymScarletI decreased. Thus, measuring ymNeongreen and ymScarletI ratio at high Cu^2+^ concentrations extended the biosensor’s linear response to 10^-3^ M Cu^2+^. Interestingly, the declining ymScarletI signal suggested a downregulation of the plasmid copy number, probably because the cells needed to curb expression from the strongly induced promoter.

We also looked at the kinetics of the promoter induction, which also illuminated nuances of the *CUP1* regulation. Flow cytometry showed symmetrical unimodal distribution for all four inducible promoters, with their median fluorescence increasing over time (Figure 2D). Thus, for each strain, all cells within the population became induced. The rate of promoter induction (Figure 2E) was similar among strains, except for pCu2, which carried the entire *CUP1* intergenic region and had lower basal activity but reached pCu1 within 30 min. This observation suggests that without Cu^2+^, the region proximal to *YHR054C* downregulates *CUP1*. Moreover, the *minCUP1* promoter was required for this regulation, as the effect was absent in *minCYC1*-bearing pCu3 and pCu4. Thus, all four inducible promoters were activated in the entire cell population within 30 minutes after adding Cu^2+^, with the combination of *YHR054C*-adjacent region and *minCUP1* promoter fostering faster induction and higher dynamic range.

### Overexpression of Cup2 transactivator

To increase the system’s sensitivity, we added to pCu plasmids a strong constitutive *CUP2* expression cassette driven by the *TDH3* promoter, thus obtaining the pCu-CUP2 plasmid series (Figure 3A). Compared to the pCu series, all new constructs had a higher basal ymNeongreen/ymScarletI ratio, and several had better detection limits and linear range (Figure 3B). While the detection limit of pCu1-CUP2 decreased, the detection limit of pCu2-CUP2 remained unchanged, although its linear range increased. The detection limit of pCu3-CUP2 and pCu4-CUP2 rose a hundred-fold, as did their linear range, allowing them to detect as little as 10^-8^ M Cu^2+^. Moreover, *minCYC1*-based pCu3-CUP2 and pCu4-CUP2 outperformed *minCUP1-based* constructs in dynamic range and reproducibility. Thus, plasmids pCu3-CUP2 and pCu4-CUP2 allowed for a highly sensitive and reproducible detection of copper, ranging from 10^-8^ to 10^-3^ M Cu^2+^.

**Figure 3:**
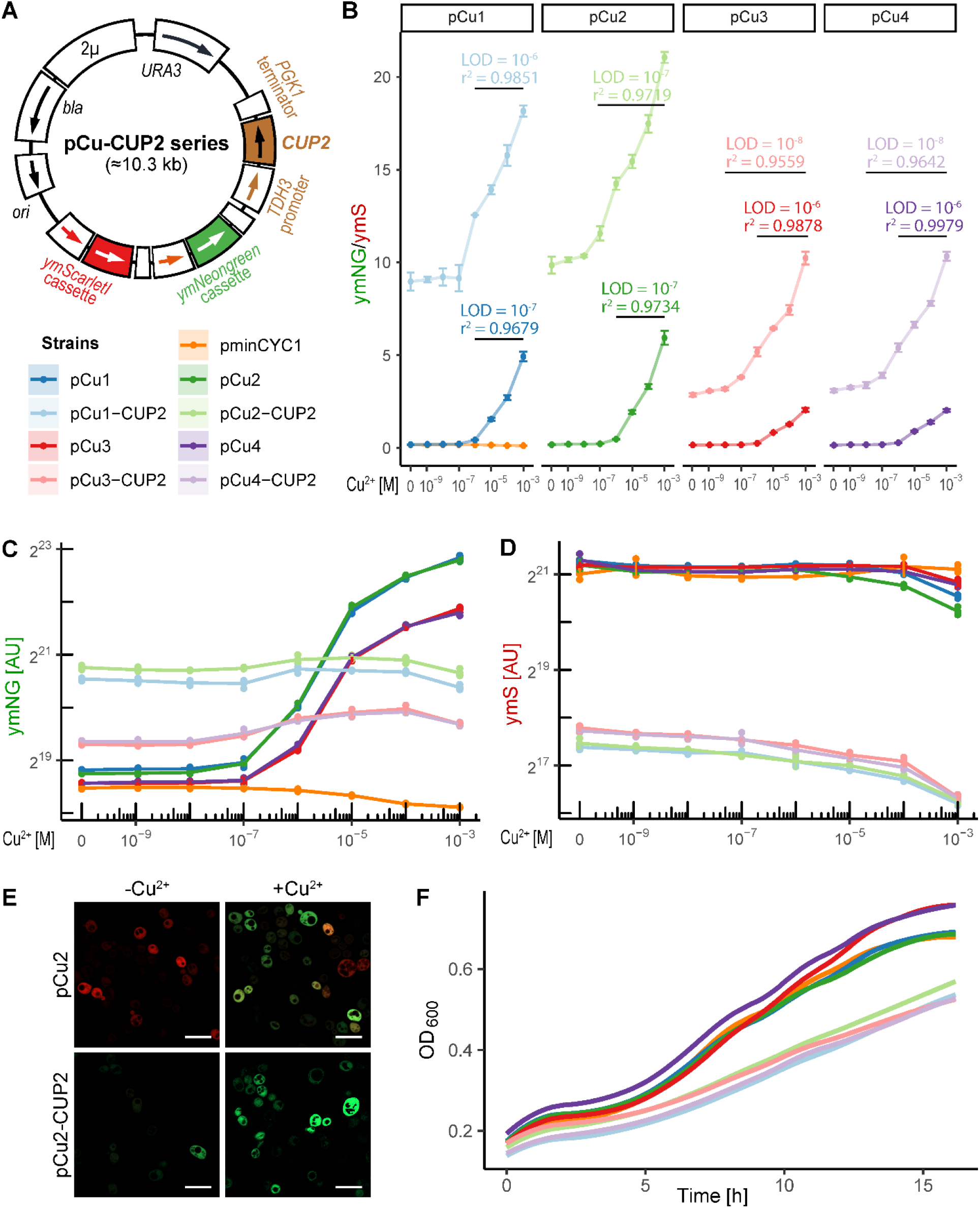
Copper biosensors overexpressing Cup2. A) Plasmids of the pCu-CUP2 series. The labels correspond to those in Figure 1B. B) Comparison of the ymNeongreen/ymScarletI ratio of copper biosensors with Cup2 overexpression (lighter colours) and without it (darker colours). C) ymNeongreen signal. D) ymScarletI signal. Other labels are as in Figure 2. E) Fluorescence micrographs of cells carrying pCu2 and pCu2-CUP2 plasmids, grown in medium without copper (-Cu^2+^) or supplemented with 100 μM copper (+Cu^2+^), expressing ymScarletI and ymNeongreen. Scale bar denotes 10.0 μm. F) Growth curves of copper biosensors.

Interestingly, the higher baseline of pCu-CUP2 plasmids did not derive from ymNeongreen, whose expression rose only two-to four-fold (Figure 3C). Instead, the effect was mainly due to a ten-fold drop in ymScarletI (Figure 3D), which was also seen in fluorescence micrographs (Figure 3E). The rise of ymNeongreen and the drop of ymScarletI were the strongest in the *minCUP1-based* pCu1-CUP2 and pCu2-CUP2, suggesting synergy between *minCUP1* promoter and Cup2 binding sites.

The increased detection limit and linear range of pCu3-CUP2 and pCu4-CUP2 stemmed from the sensitivity and accuracy of the ratiometric measurement as in these systems the rising Cu^2+^ concentration barely increased the ymNeongreen signal, while the already faint ymScarletI signal decreased. Thus, although pCu3-CUP2 and pCu4-CUP2 performed outstandingly, they never obtained the fluorescent intensity of pCu3 and pCu4. Finally, all pCu-CUP2 plasmids slowed cell growth (Figure 3F), with cells retaining normal morphology (Figure 3E) but reaching lower OD_600_ values in the stationary phase (data not shown). Thus, pCu3-CUP2 and pCu4-CUP2 can detect copper concentrations as low as 10^-8^ M but interfere with yeast growth, and so pCu1 and pCu2 might be more convenient when such sensitivity is not required.

### Partial deletions of Cup2 transactivator

We wondered if we could fine-tune the system further, boosting the growth and fluorescent signal of pCu4-CUP2 cells while retaining their extreme Cu^2+^ sensitivity. As overexpressed transactivators can slow growth through overly strong activation domains (Gill and Ptashne, 1988; Meyer et al., 1989), we tried to pinpoint the Cup2 activation domain. Alphafold’s prediction of Cup2 structure (Figure 4A, Uniprot Accession P15315) showed that the transactivator’s DNA-binding and regulatory regions, residing in the N-terminal half of the protein, were well-structured. On the other hand, the C-terminal half of the protein appeared intrinsically disordered (Ruff and Pappu, 2021), which is characteristic of activation domains (Ravarani et al., 2018). Moreover, a localised search for activating sequences identified a 75% match to the 9aaTAD pattern (Piskacek et al., 2016) in the Cup2 211-219 residues that fold into solitary alpha-helix at protein’s C-terminus.

**Figure 4:**
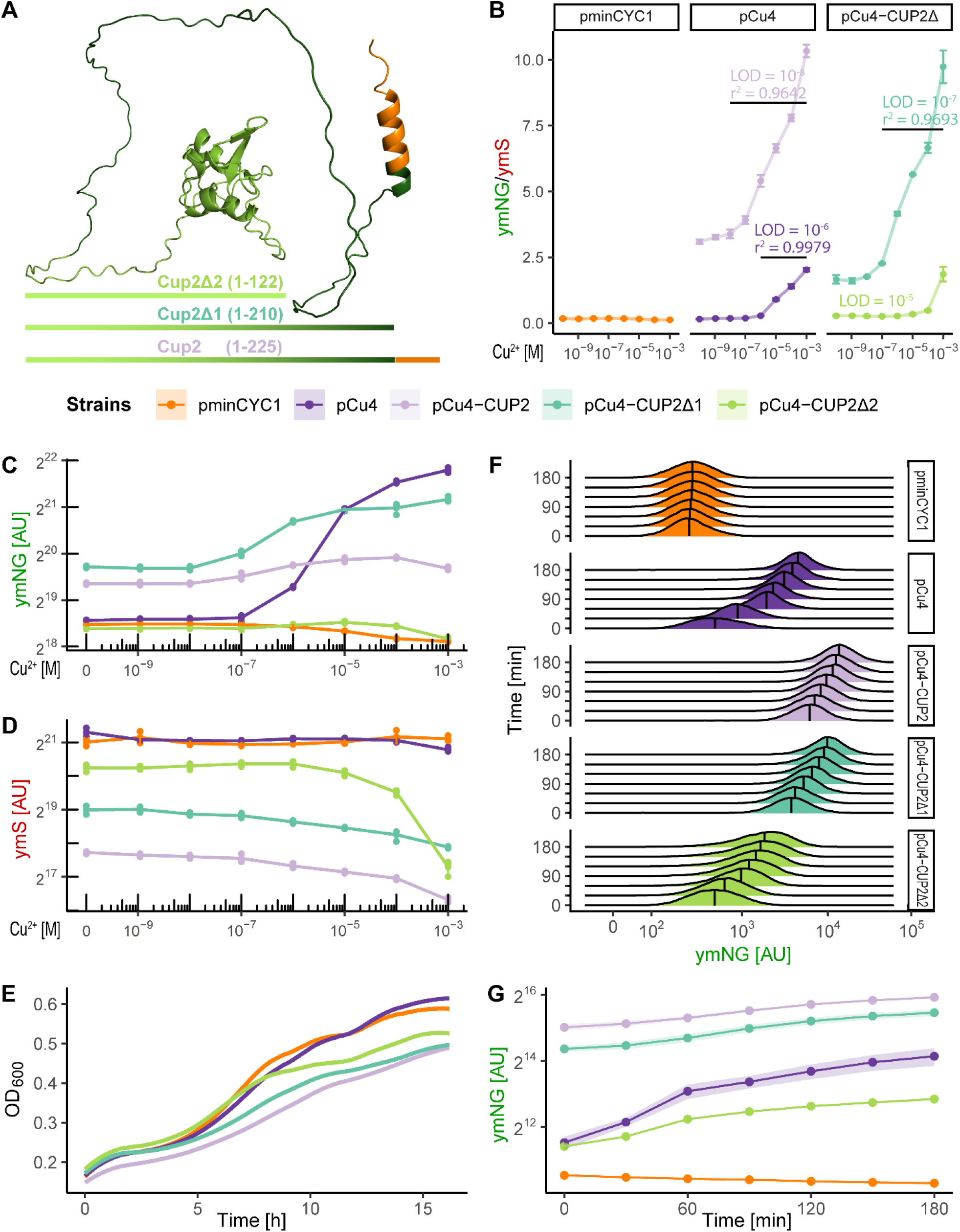
Copper biosensors overexpressing truncated Cup2. A) Alphafold-predicted structure of Cup2 and its trimmed variants. Cup2Δ1 contains both light-green and dark-green regions of Cup2, but not the orange region, while Cup2Δ2 contains only the light-green region. Numbers in parentheses denote amino acid residues contained within each variant. B) The ymNeongreen/ymScarletI ratio for control construct, pCu4 biosensor without and with Cup2 overexpression, as well as biosensors overexpressing trimmed Cup2 variants. C) ymNeongreen signal. D) ymScarletI signal. Other labels are as in Figure 2. E) Growth curves. F) The distribution of ymNeongreen signal in cells induced with 100 μM Cu^2+^, measured by flow cytometry. The ymNeongreen cassette is integrated into the chromosome. Vertical lines within distributions denote median fluorescence. G) Kinetics of promoter induction with 100 μM Cu^2+^ in cells with ymNeongreen cassette integrated into the chromosome, measured by flow cytometry.

Based on structural information, we tried to dampen the activity of plasmid-based Cup2. Thus, we constructed two truncated Cup2 variants, lacking 15 or 103 C-terminal amino acid residues (Cup2Δ1 and Cup2Δ2, respectively; Figure 4A). The constructs’ properties differed markedly. Compared to pCu4-CUP2, plasmid pCu4-CUP2Δ1 imparted poorer detection limit and linear range but increased dynamic range (Figure 4B) and strengthened ymNeongreen and ymScarletI signals (Figures 4C and 4D). On the other hand, plasmid pCu4-CUP2Δ2 effectively repressed the promoter, lowering its detection limit and ymNeongreen signal below those of pCu4 (Figures 4B and 4C), thus suggesting that Cup2Δ2 competitively inhibits DNA binding of the endogenous, chromosomally encoded Cup2. Such inhibition also interfered with regular copper-detoxifying Cup1 response, thus undermining growth at higher copper concentrations (Figure 4D). Interestingly, although pCu4-CUP2, pCu4-CUP2Δ1, and pCu4-CUP2Δ2 carried the same promoters, they differed in ymScarletI signals, suggesting that plasmid copy number also depended on the protein that plasmid expressed. Finally, compared to pCu4-CUP2, plasmid pCu4-CUP2Δ1 had no impact on the rate of growth (Figure 4E) and promoter induction (Figures 4F and 4G), while pCu4-CUP2Δ2 grew faster than pCu4-CUP2 but slower than pCu4 (Figure 4E). As such, growth curves also imply that cells grew slowly due to overly strong activation domains. Overall, truncated Cup2 variants performed worse than full-length Cup2, suggesting that potent activation domains slow growth and weaken the fluorescent signal but significantly increase detection limit and linear range.

### Cup2 variants with heterologous activation domains

As previous results suggested that a more potent activation domain benefits the biosensor, we fused truncated Cup2 variants with strong heterologous activation domains. Thus, to both Cup2Δ1 and Cup2Δ2, we appended activation domains VP64 from herpes simplex virus (Beerli et al., 1998), EDLL2 from *Arabidopsis thaliana* (Naseri et al., 2017), or their combination.

Hybrid Cup2 transactivators underlined the importance of balanced activation. Fusing Cup2Δ1 with either VP64 or EDLL2 lowered the systems’ detection limit ten-fold, and linking both domains to it lowered it even more (Figure 5A). On the contrary, fusion of Cup2Δ2 with either domain was beneficial, with construct carrying linked domains rivalling the non-truncated Cup2. The ymNeongreen and ymScarletI also reflected variation in construct performance (Figures 5B, 5C, and 5D, 5E, respectively), with the expression of ymScarletI closely matching the growth rates (Figures 5F and 5G). Thus, adding domains to mostly intact Cup2 creates an overpowering transactivator, harming the system. In contrast, adding the same domains to the solitary DNA-binding domain of Cup2 is beneficial.

**Figure 5:**
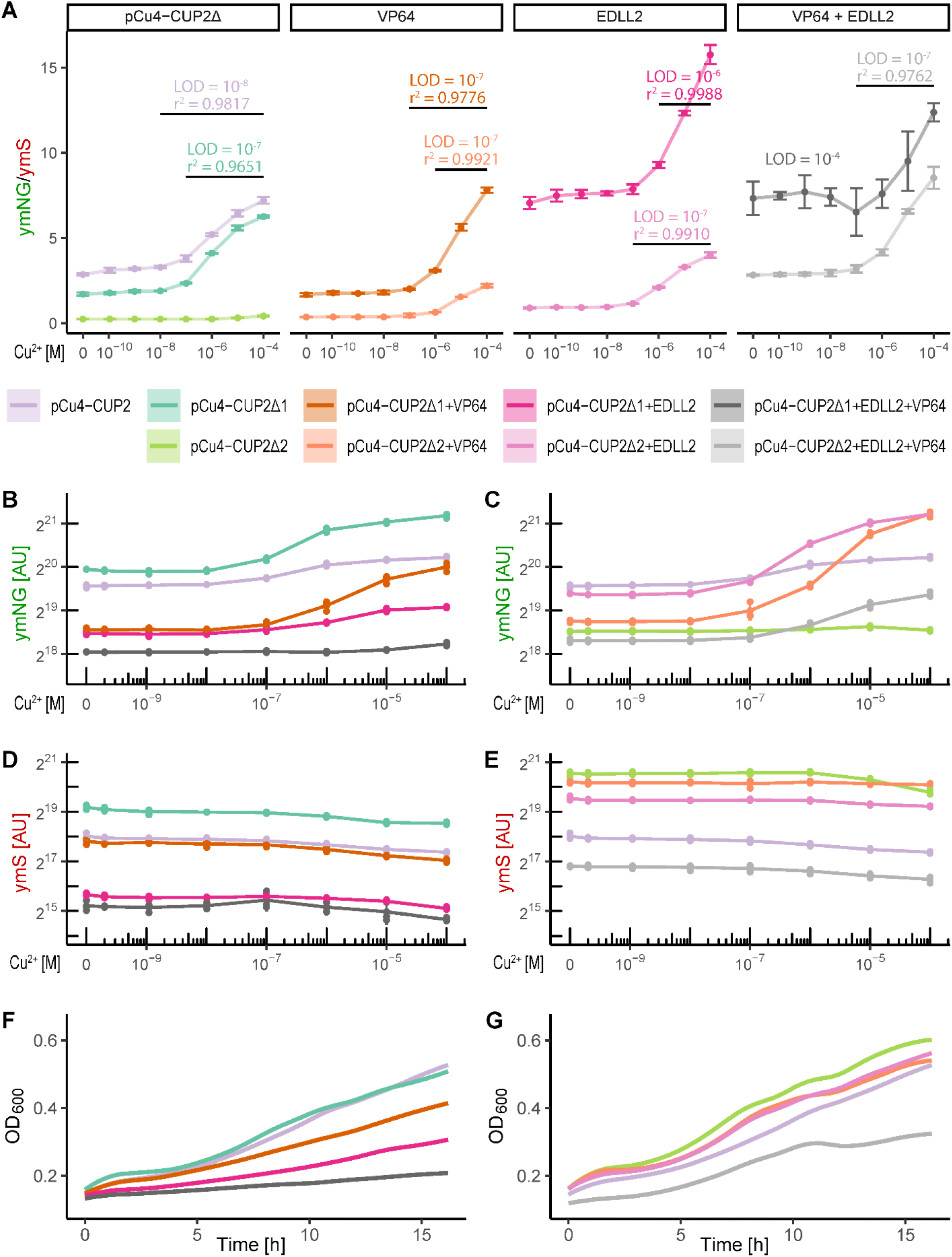
Copper biosensors overexpressing Cup2 with different activation domains. A) Comparison of the ymNeongreen/ymScarletI ratio for pCu4 biosensor overexpressing Cup2, its trimmed variants, and combination of trimmed variants with VP64 and EDLL2 activation domains. B) ymNeongreen signal of Cup2Δ1-based variants. C) ymNeongreen signal of Cup2Δ2-based variants. D) ymScarletI signal of Cup2Δ1-based variants. E) ymScarletI signal of Cup2Δ2-based variants. F) Growth curves of Cup2Δ1-based variants. G) Growth curves of Cup2Δ2-based variants.

### Biosensor specificity and comparison with other whole-cell copper biosensors

As Cup2 binds Ag^+^ *in vitro* (Buchman et al., 1989), we determined the specificity of our system by trying to detect related metals in concentrations up to 10^-3^ M. However, we observed no response when the medium was supplemented with Co^2+^, Ni^2+^, Zn^2+^, Pa^2+^, Cd^2+^, Ag^+^, or Au^+^ (data not shown). Thus, our system is highly specific for Cu^2+^ ions.

Finally, we compared pCu4-CUP2, our best performing biosensor, with twelve others reported in the literature (Figure 6). This comparison showed that pCu4-CUP2 outperformed other eukaryotic whole-cell biosensors, having at least a hundred-fold wider linear range and at least ten-fold better detection limit. While the state-of-the-art prokaryotic *E. coli*-based biosensors performed better than eukaryotic ones, only one had detection limit comparable to pCu4-CUP2 and none had as wide linear range. Thus, the system developed in this work distinguishes itself by an outstanding combination of specificity, detection limit, and linear range.

**Figure 6:**
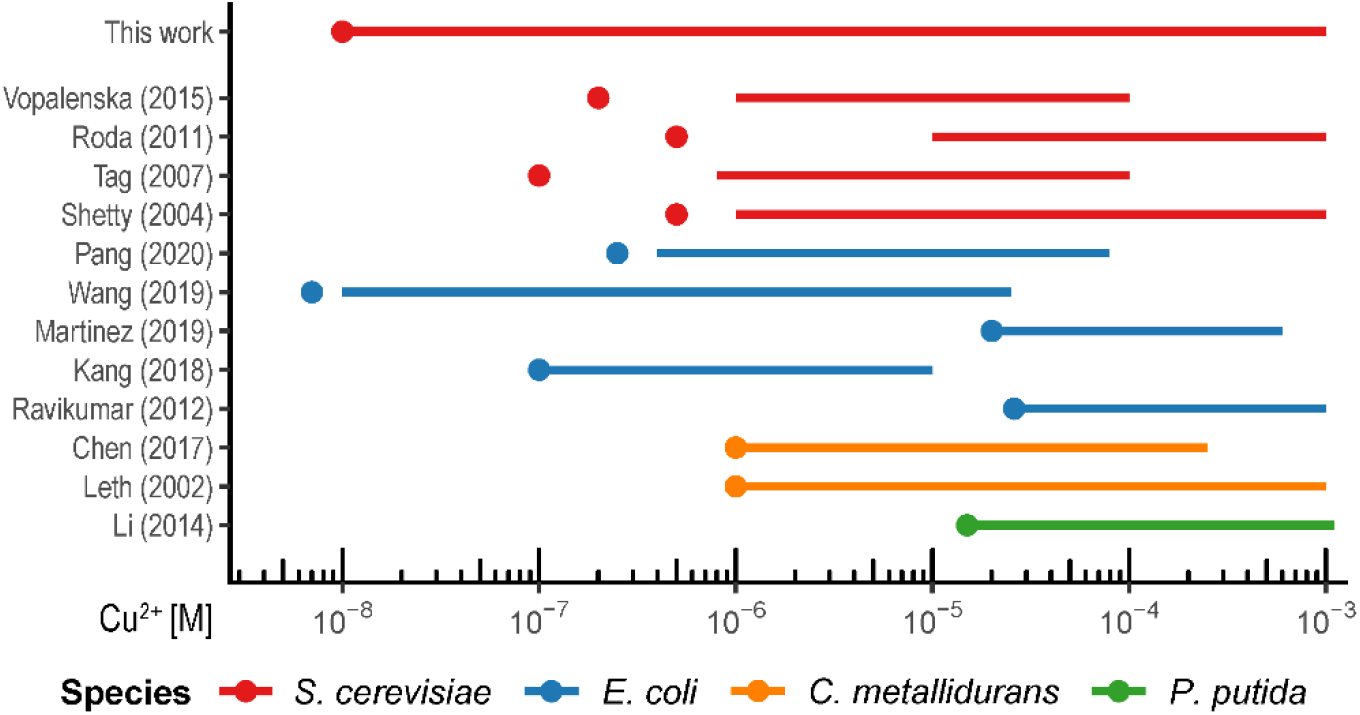
Comparison of pCu4-CUP2 with other published copper biosensors. The horizontal line marks the linear range and the point next to it marks the detection limit, with colours denoting different species.

## Discussion

In this work, we redesigned yeast *S. cerevisiae* into a whole-eukaryotic-cell dual-reporter biosensor able to detect 10^-8^ to 10^-3^ M of bioavailable copper. For this purpose, we constructed two native and two hybrid variants of the *CUP1* promoter and tested them in wild-type cells and in cells overexpressing Cup2 transactivator. While the native variants performed better in wild-type cells, they were surpassed by hybrid variants in cells overexpressing Cup2 transactivator. However, the abundance of Cup2 slowed down cell growth, prompting us to engineer its activation domain. While trimming this domain helped growth but worsened sensitivity, strengthening it worsened both metrics, suggesting further development should rely on subtler domain modifications.

Biosensors developed in this work can detect a concentration of bioavailable copper as low as 10 nM and have extremely wide linear ranges, from 0.01 μM to 1000 μM (5 log scale), thus ranking amongst the most sensitive whole-cell copper biosensors. Moreover, they measure only copper capable of entering the eukaryotic cell, disregarding non-bioavailable copper and other heavy metals. Most other whole-cell biosensors are prokaryotic, which are less sensitive and have a narrower linear range. However, they still perform better than existing eukaryotic systems, primarily constructed in yeast *S. cerevisiae*. On the other hand, voltammetry and amperometry-based eukaryotic biosensors, incorporating lyophilised or non-living cells in the carbon paste electrode, regularly detect 0.05 μM Cu^2+^ (Alpat et al., 2008; Alpat et al., 2007; Yüce et al., 2010) but work by measuring the adsorption of metal ions to the cell wall, and so their specificity is uncertain. Thus, the system described in this work distinguishes itself through high specificity, low detection limit, and wide linear range.

The originality of this work also relies on the design of a novel ratiometric, i.e. dual-reporter whole-cell biosensor, which is one of the only few such systems currently described (Mirasoli et al., 2002; Roda et al., 2011). We used optimised fluorescent proteins as they are stable in yeast cells and detectable in small amounts without cell lysis or external substrate. With ymScarletI as an internal reference system, we evaded OD_600_, an unreliable measure of cell density in polluted and toxic environments, such as those contaminated with a mixture of heavy metals. The ymScarletI also monitored viability, showing that cells were alive, even when they detected no copper. Interestingly, although rarely used in whole-cell biosensors, ratiometric systems are regularly employed for *in vivo* measurements of intracellular metabolites as FRET biosensors (Bhuckory et al., 2019) or circularly-permuted fluorescent proteins (Kostyuk et al., 2019; Liu et al., 2020). Thus, the presented ratiometric system has clear advantages over single reporters, extending biosensor’s detection limit and linear range, and serving as a dual-sensing reporter for Cu^2+^ detection and cell viability.

We constructed and compared four versions of the *CUP1* promoter without heat shock element (HSE), ensuring it reacts only to Cu^2+^ and not high temperature or oxidative stress (Hottiger et al., 1994; Tamai et al., 1994), thus making it more independent of the cell’s physiological status. While others used this promoter, they ignored HSE and employed only the Cup2-binding region and *minCUP1* promoter, discarding the *YHR054C*-adjacent sequence. We showed that the *YHR054C*-adjacent sequence cooperates with *minCUP1* promoter to affect induction kinetics and that the *minCUP1* promoter considerably affects the biosensor’s detection limit and dynamic range, although it does not carry Cup2-binding sites. Interestingly, Shen et al. (2001) showed that Cup2 binding repositions nucleosomes over the entire *CUP1* gene, while Badi and Barberis (2002) noticed that the region upstream of *CUP1* UAS restores impaired activation of *CUP1* transcription, even suggesting that the *YHR054C* sequence itself has a regulatory role. Moreover, overexpression of Cup2 benefited the *minCYC1*-based biosensors but not the *minCUP1-based* biosensors, suggesting an interaction between the Cup2-binding region and *minCUP1* promoter, which is disrupted in hybrid promoter but restored at higher Cup2 concentration. Thus, the *CUP1* promoter seems to have evolved as one compact, well-tuned transcription unit, with both Cup2-binding and non-binding segments working together to regulate transcription.

The Cup2 overexpression enhanced under-performing *minCYC1*-based constructs by improving their detection limit, operational, and dynamic range, even beyond those of *minCUP1* constructs. This result highlighted difficulties in predicting construct performance and underlined the importance of testing many variants. However, overexpression of Cup2 lowered ymNeongreen and ymScarletI signals, complicating their detection and reducing reproducibility, with the overexpression system ultimately performing well because of the dual-reporter ratiometric approach.

The Cup2 overexpression also hampered growth, probably through squelching (Schmidt et al., 2016), i.e. cell-wide disturbance of transcription caused when overexpressed transactivator binds up too much Mediator, causing its cell-wide shortage. In support of this hypothesis, Sanborn et al. (2021) recently demonstrated that the Cup2 activation domain directly binds Mediator, as do most yeast transcription factors. Another possibility is that the internally disordered C-terminal domain aggregates and even phase-separates at higher concentrations (Becskei, 2020). In either case, we restored growth by trimming C-terminal half of Cup2 but thus created a transcriptional repressor that competes with chromosomally encoded Cup2 for binding sites, blocking copper response. These results suggest that similar overexpression of only DNA-binding domain could serve as a general strategy for damping unwanted transcriptional responses.

We further adjusted the biosensor by building upon or exchanging native Cup2 activation domain with foreign sequences, e.g. viral VP64 or plant EDLL2 domains. We noted that strong activation domains slowed growth, especially when present together, paralleling the observation that domains present *in tandem* bind Mediator stronger than expected due to synergistic effect of their interaction sites (Sanborn et al., 2021). Curiously, this effect could be employed to limit the growth rate, and thus harnessed for biocontainment and biosafety (Wang and Zhang, 2019).

We constructed and compared a total of 16 copper biosensors, which is an unusual approach given that most publications characterise only one variant. However, such an approach allowed us to deduce some design guidelines and inspired further modifications. For example, while we focused on exchanging entire activation domains, an alternative approach would be to change single amino acids, as such mutations can have a striking effect (Sanborn et al., 2021), while allowing for a more precise tuning of the signal. Moreover, the detection limit might be improved by deleting an array of *CUP1* genes, which down-regulate their own expression (Wright et al., 1988) and are over-induced in strains overexpressing Cup2 (Huibregtse et al., 1989). Additional strategies could rely on modifying chromosomally encoded *CUP2* or introducing more Cup2 binding elements into the inducible promoter.

## Conclusion

In this work, we took advantage of the native copper response of yeast *S. cerevisiae* and redesigned it. We constructed the next generation eukaryotic whole-cell copper biosensor exhibiting very high sensitivity, specificity, and wide linear range, allowing detection of bioavailable cooper from 10^-8^ to 10^-3^ M. For this purpose, we combined an unprecedented dual-reporter strategy with significant promoter and transactivator engineering. Our approach points out a new paradigm for developing and improving any eukaryotic whole-cell biosensor.

## Acknowledgements

We thank all members of Bénédetti/Vallée group, and especially Michel Doudeau and Fabienne Godin for technical support. We also thank David Gosset for his help with flow cytometry and fluorescence microscopy.

## CRediT author statement

**Bojan Žunar**: Conceptualization, Methodology, Software, Validation, Investigation, Formal analysis, Visualization, Writing - Original Draft, Writing - Review & Editing. **Christine Mosrin**: Resources, Writing - Review & Editing. **Héléne Bénédetti**: Resources, Supervision, Writing - Review & Editing. **Béatrice Vallée**: Conceptualization, Methodology, Resources, Writing - Review & Editing, Supervision, Project administration, Funding acquisition.

